# PathMe: Merging and exploring mechanistic pathway knowledge

**DOI:** 10.1101/451625

**Authors:** Daniel Domingo-Fernández, Sarah Mubeen, Josep Marín-Llaó, Charles Tapley Hoyt, Martin Hofmann-Apitius

## Abstract

**Background:** The complexity of representing biological systems is compounded by an ever-expanding body of knowledge emerging from multi-omics experiments. A number of pathway databases have facilitated pathway-centric approaches that assist in the interpretation of molecular signatures yielded by these experiments. However, the lack of interoperability between pathway databases has hindered the ability to harmonize these resources and to exploit their consolidated knowledge. Such a unification of pathway knowledge is imperative in enhancing the comprehension and modeling of biological abstractions.

**Results:** Here, we present PathMe, a Python package that transforms pathway knowledge from three major pathway databases into a unified abstraction using Biological Expression Language as the pivotal, integrative schema. PathMe is complemented by a novel web application (freely available at https://pathme.scai.fraunhofer.de/) which allows users to comprehensively explore pathway crosstalks and compare areas of consensus and discrepancies.

**Conclusions:** This work has harmonized three major pathway databases and transformed them into a unified schema in order to gain a holistic picture of pathway knowledge. We demonstrate the utility of the PathMe framework in: i) integrating pathway landscapes at the database level, ii) comparing the degree of consensus at the pathway level, and iii) exploring pathway crosstalk and investigating consensus at the molecular level.

## 1. Background

The interpretations of molecular signatures that are typically yielded by genome-scale experiments are often supported by pathway-centric approaches through which mechanistic insights can be gained by pointing at a set of biological processes. Thus, parallel to the development of novel data-driven approaches, pathway databases emerged as comprehensive resources that could be used to complement analyses with prior knowledge. These resources have embraced standard file formats and schemata in order to facilitate the exchange of pathway knowledge. However, each resource has chosen a different one and though these formats possess overlapping capabilities to produce computational models of biology, their intended purposes and applications are somewhat distinct. For instance, Systems Biology Markup Language (SBML) is a standard for the representation of computational models of systems biology, Systems Biology Graphical Notation (SBGN) facilitates the storage and exchange of signaling pathways, metabolic networks and gene regulatory network information, and Biological Pathway Exchange (BioPAX) has been designed with the purpose of establishing a common exchange format for biological pathway data (Hucka *et al.*, 2003; Demir *et al.*, 2010; Le Novere *et al.*, 2009). A variety of formats offer the scientific community multiple approaches to curate pathway knowledge. However, a multitude of diverse formats and a lack of interoperability between them tends to hamper efforts to collate the knowledge contained in pathway databases. In practice, this has led to the generation of data silos derived from the gradual detachment of complementary work from different research groups which use distinct modeling languages. Therefore, metadatabases such as Pathway Commons (Cerami *et al.*, 2011) and ConsensusPathDB (Kamburov *et al*., 2008), which incorporate several different primary resources in their data warehouses, and integrative software applications such as *graphite* (Sales *et al*., 2018) and OmniPath (Türei *et al*., 2016) have been created with the primary intention to integrate pathway knowledge from multiple databases. Beyond these, other approaches such as those taken in PathCards (Belenky *et al*., 2015), RaMP (Zhang *et al*., 2018), and ComPath (Domingo-Fernández *et al.*, 2019), have focused on integrating gene sets and chemical knowledge related to pathways, but without including their topological information (i.e, relationships were excluded from the network). For instance, ComPath, the precursor of this work, harmonized pathway information at the gene level in order to conduct extensive manual curation that mapped and cross-referenced pathway representations across databases. This mapping catalog reveals which pathways are covered by which database (e.g., pathway in resource X is equivalent to pathway in resource Y) and facilitates comparing the results of pathway enrichment methods.

The representation of pathway knowledge can span across several scales including molecular events, cellular processes and/or phenotypes, which are captured in varying degrees by integrative resources. For example, ConsensusPathDB and *graphite* effectively account for and harmonize metabolites, genes, and proteins when integrating pathways from multiple databases, but exclude biological types at higher order scales, such as biological processes, and other entities, such as miRNAs. On the other hand, Pathway Commons can incorporate multiple scales of biology by retaining original entity identifiers; however, it does not directly harmonize biological entities.

An ongoing challenge in harmonizing pathway resources is the use of distinct nomenclatures by individual databases. For example, for gene and gene products there exist several standard terminologies such as ENTREZ (Maglott *et al.* 2005), UniProt (Apweiler *et al.*, 2004), Ensembl (Hubbard *et al.*, 2002), and HGNC (Povey *et al.*, 2001), or for chemicals, ChEBI (Hastings *et al., 2015*), ChEMBL (Gaulton *et al.*, 2011), and PubChem (Bolton *et al.*, 2008). Despite the availability of standard terminologies, some resources still assign biological entities and concepts to internal database identifiers. Therefore, mappers are necessary to normalize identifiers and facilitate resource harmonization (van Iersel *et al.*, 2010). Similarly, the harmonization of biological relationships is required to unify heterogenous networks. While several format translators can convert interactions across formats (Bohler *et al.*, 2016; Bonnet *et al.*, 2013; Demir *et al*., 2013; Gyori *et al.*, 2017; Wrzodek et *al.*, 2011), the process of harmonizing relationships, or edges in pathway networks, is not trivial; thus, hampering an integrative approach comprising several databases.

Just as pathway databases should be regularly updated to incorporate continual changes in pathway definitions, pathway metadatabases should also be updated in parallel to reflect such changes; it has been shown that by using outdated resources, results of studies are strongly influenced, and follow-up studies are negatively impacted (Wadi *et al.*, 2016). Correspondingly, approaches to harmonize pathway data also require these considerations or they too would be subject to similar liabilities. Moreover, pathway analysis software have been recently complemented with user-friendly exploratory tools and applications such as Pathway Commons, PathVisio (Kutmon *et al*., 2015), Cytoscape.js (Franz *et al*., 2015), or NDEx (Pratt *et al*., 2015), which have been specifically designed for the visualization of individuals pathways and biological networks, including at a finer, more granular level. While the scientific community has greatly benefited from the development of these tools, there is still the need to develop applications that focus on visualizing the consensus and crosstalk between multiple, disparate pathway representations. While previously mentioned attempts have succeeded in accumulating and increasing the availability of database content, there has not yet been a systematic evaluation that investigates the degree of overlap or the amount of agreements/discrepancies in related or equivalent pathways from different databases. Previous comprehensive comparisons of database content were restricted to single or small sets of pathways because of the considerable amount of manual intervention (e.g., entity/relationship normalization, image reconstruction, etc.) required to shed light on the degree of overlap of equivalent pathways (Stobbe *et al.*, 2011; Chowdhury and Sarkar, 2015). Conversely, conducting a systematic comparison requires harmonization of entities and biological interactions across databases and minimizing pathway information loss whilst accommodating databases into an interoperable schema (i.e., retain most of the different biological abstractions that each database offers in the transformation process). Finally, connecting and integrating pathway knowledge can enhance pathway enrichment analyses, as has already been demonstrated in a more simplistic approach by Minadakis *et al*., as well as drive curation and new experimentation by highlighting the consensus, discrepancies, and unexplored areas of the pathway landscape.

Here, we introduce PathMe, an extensible package that harmonizes multiple databases using Biological Expression Language (BEL) as a common interoperable schema and enables pathway knowledge evaluation and exploration powered by a stand-alone web application with a special focus on highlighting pathway crosstalks and consensus.

## 2. Implementation

PathMe framework is comprised of two parts: the open-source Python package that converts the different database formats into BEL and the web application that allows for the exploration of the resulting networks **(Figure 1)**.

**Figure 1.**
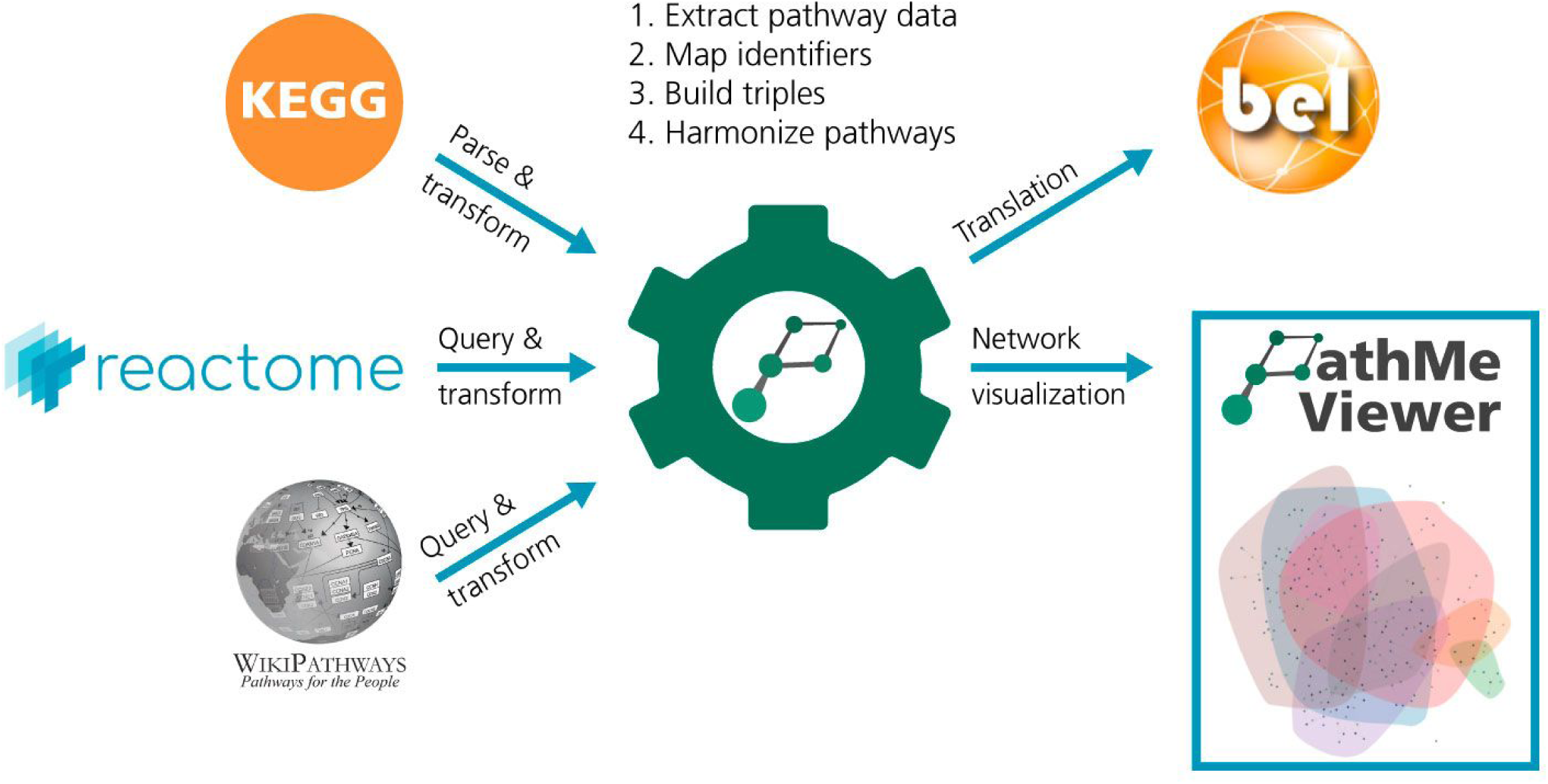
Design of the PathMe framework. The PathMe software package facilitates the transformation of pathway content into BEL. The initial step consists of extracting, parsing, and/or querying content from each pathway database to retrieve entities, concepts, interactions and reactions, and their associated metadata. Subsequently, database specific identifiers for all entities are unified to stable and consistent ones, where possible. Data are then directly mapped into equivalent BEL nodes and edges, translating all human pathways from the databases into BEL. Finally, an interactive pathway viewer is implemented such that any combination of pathways, represented as BEL networks, can be explored and the consensus surrounding pathway knowledge can be directly compared.

### 2.1. The PathMe Python package

#### 2.1.1. Integrating knowledge across pathway databases

Integrating pathway knowledge from multiple databases first requires transforming the content of each database into a common underlying schema. While multiple triple-based formats can be used to formalize pathways in system biology, we adopted BEL as the pivotal unifying schema since it provides a reasonable trade-off between expressivity and standardized organization. Until now, we have implemented parsers for three major databases (i.e., KEGG, Reactome, and WikiPathways (Kanehisa *et al*., 2016; Fabregat *et al.*, 2018; Slenter *et al.*, 2017)) that extract pathway information and serialize it to BEL. As the principal goals of PathMe are to enable direct comparisons and explorations of pathways from different databases, cross-database mappings of identifiers and relation types are required. Accordingly, the parsers harmonize molecular entities to identifiers from standard nomenclatures as well as interaction types into their corresponding BEL relationships.

In order to harmonize entities, we prioritized standard nomenclatures for each of the modalities (e.g., genes, proteins, metabolites, etc.) included in the three studied databases (**Tables S1**, **S4**, **and S6 in Supplementary Text)**. HGNC was the top-level priority namespace for genes and gene products (Povey *et al.*, 2001). HGNC was selected as it is recognized as an authority for standard nomenclature assignments and annotations for human genes and because the software is primarily concerned with converting human pathways. In the absence of HGNC identifiers, lower level priority namespaces were used to derive the top level HGNC identifier assignment. For instance, we aimed to use intermediate level UniProt identifiers (Apweiler *et al.*, 2004) to map back to HGNC identifiers. If mappings to the prioritized namespaces were not available, genes and gene products retained their database-specific identifiers and were assigned to namespaces designated by their respective databases in order to maximize the retrieval of entities from each resource. Similarly, metabolites were prioritized to preferentially obtain ChEBI identifiers because of ChEBI’s wide usage as a source of manually curated stable identifiers and annotations for small chemical compounds (Hastings *et al., 2015*). In their absence, either PubChem identifiers were assigned or, once again, they retained their database-specific identifiers. Once entities were assigned to standardized identifiers, the modalities defined by the source databases were mapped to their corresponding BEL node classes (e.g., gene, protein, metabolite, biological process, etc.). Efforts were made to accommodate entities not readily mappable to BEL nodes by using BEL node classes which can incorporate flexibility in their definitions. For instance, unspecified physical entities in WikiPathways are given the abstract class label, ‘DataNode’; these ‘DataNodes’ were mapped to BEL abundances, a category that represents the abundance of a biological entity such as a chemical or an unspecified molecule. Entity class mappings from the source databases to BEL are summarized in **Tables S2**, **S5**, **and S7 of the Supplementary Text.**

Similar to the normalization of biological entities into a standardized nomenclature and their translation into corresponding BEL entity classes, distinct relationships utilized in the biological networks of different databases must too be normalized. While the versatility of BEL permitted the successful transformation of all relationships from Reactome and WikiPathways, four KEGG relationships (i.e., hidden compound, state change, dissociation, and missing interaction) could not be translated into BEL due to the lack of correspondingly equivalent edges in the BEL syntax. However, these four relationships represent non-causal interactions between biological entities and are also minimally utilized by KEGG curators. Mappings between edges from the source databases to BEL are reported in **Supplementary Tables S3**, **S5 and**, **S7.**

#### 2.1.2. Implementation details

PathMe relies on the individual parsers that convert the original formats from the databases to BEL. Each parser is implemented using libraries that enable the manipulation and transformation of its corresponding schemata (i.e., RDFLib for Resource Description Framework (RDF) and the *xml* Python package for Extensible Markup Language (XML)). Moreover, the parsers are structured into their own packages inside the main Python module to facilitate the inclusion of additional database parsers in the future. During the entity normalization process, mappings across identifiers are facilitated through the numerous packages included in the Bio2BEL framework (https://github.com/bio2bel) (**Table S9 in the Supplementary text**). After the normalization, entities and their relationships in each pathway are translated to BEL using the internal domain specific language (DSL) and the *BELGraph* class of the PyBEL Python software package (Hoyt *et al.*, 2017). PathMe benefits from the numerous modules implemented in the PyBEL ecosystem since it offers a variety of functionalities and algorithms that enable querying, transforming, and analyzing biological networks, as well as an export module that can output multiple formats. Finally, PathMe is distributed as a Python package through Python Package Index (PyPI) and its source code is available in GitHub at https://github.com/PathwayMerger/PathMe.

### 2.2. PathMe Viewer

#### 2.2.1. A web application to explore pathway knowledge

As discussed in the introduction, several visualization tools have focused on the exploration of biological networks, but have never attempted to study or evaluate the coverage, consensus, and crosstalks across heterogeneous networks. Since the particular use case of this work called for customized solutions (e.g., delineating boundaries or highlighting agreements when multiple pathways are being visualized), we also implemented a novel tool called PathMe Viewer to fulfill these unmet needs and complement the PathMe package. Since the target audience for this application are pathway curators and researchers, we opted to implement the viewer in the form of a user-friendly web application compatible with any device. The front-end extends the visualizations from BEL Commons (Hoyt *et al*., 2018) and provides an intuitive and interactive interface for visualizing and exploring the knowledge comprised in the pathway landscape. Moreover, the web application is complemented with analysis modules and network algorithms to query pathways or calculate their similarity as well as exporting options to multiple standard formats such as BEL, GraphML, or JSON so networks can be used in other software designed for visualization purposes such as Cytoscape (Franz *et al*., 2015) or advanced algorithmic analyses such as SPIA (Tarca *et al*., 2008). Finally, the network visualization is a stand-alone component within the web application and it remains agnostic to BEL by rendering the graphics using the Node-Link JSON data format, a standard format used by popular visualization libraries; thus, facilitating the reusability of the component out of the BEL community.

#### 2.2.2. Implementation details

PathMe Viewer follows a model-view-controller (MVC) software architecture. While the back-end is implemented in Python using the Flask microframework, the front-end is implemented in JavaScript using libraries such as jQuery (https://jquery.com), D3.js (https://d3js.org), and Bootstrap (https://getbootstrap.com). The source code is available at https://github.com/PathwayMerger/PathMe-Viewer so that all visualizations and components can be reused or extended by future applications. Furthermore, the web application is distributed through PyPI and can also be deployed with Docker which facilitates the reproducibility of this work since Docker’s automated deployment process ensures that every single instance runs with the exact same settings, regardless of the host machine. Documentation is included in the GitHub repository and it is also accessible through Read the Docs (https://pathme-viewer.readthedocs.io/en/latest/). Finally, we provide access to a public deployment of the PathMe Viewer at https://pathme.scai.fraunhofer.de.

### 2.3. Calculating pathway similarity

As an application of the software, we conducted the following protocol to evaluate the degree of overlap between the three representations of each equivalent pathway (*Case scenario II*). We used a variation of the Szymkiewicz–Simpson/Overlap coefficient **(Equation 1)**, calculated for common molecular nodes shared between the networks. To calculate a pathway similarity index, we summed the three coefficients obtained for each individual pairwise comparison and divided this number by three to normalize to a zero-to-one scale. In other words, each pathway similarity index corresponds to a normalized sum of the individual overlaps between: i) the KEGG and Reactome representation, ii) the KEGG and WikiPathways representations, and iii) the Reactome and WikiPathways representations. Therefore, the pathway similarity index (S) lies between 0 ≤ S ≤ 1 (with 0 corresponding to no overlap between any of the three sets, and 1 corresponding to three fully overlapping sets).

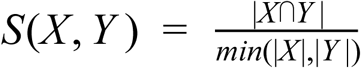

**Equation 1.** The Szymkiewicz-Simpson coefficient calculates the similarity between two sets (X and Y) where 0 ≤ S ≤ 1. The similarity is the size of the intersection of the two sets divided by the size of the smaller set. In this case, the sets correspond to the number of individual molecular entities excluding group nodes in the BEL graph, this is discussed in detail in the **Supplementary Text**.

## 3. Results

In sections 3.1 and 3.2, we present the main functionalities of the PathMe software and web application respectively, while section 3.3 outlines the architecture and design of the framework. Next, sections 3.4, 3.5, and 3.6 present three case scenarios applied at increasingly granular scales of pathway knowledge to illustrate the usability of the framework in database integration from a global, database-wide perspective to a detailed, pathway level one.

### 3.1. PathMe *functions*

The PathMe package offers a set of functionalities for the set of databases incorporated thus far: i) download the raw pathway files, ii) generate BEL networks and export them as binary data, iii) summarize the transformed content, and iv) calculate detailed network statistics (e.g., number of nodes, edges and their types) **(Table 1)**. Moreover, database specific features include functionalities to flatten all group nodes (e.g., protein complexes, gene families, etc.) in KEGG and exclusively parse canonical pathways from WikiPathways and Reactome. In conclusion, these functionalities combined with the ones already offered by the PyBEL ecosystem assist bioinformaticians in transforming, exploring, and analyzing the generated pathway networks.

**Table 1.**
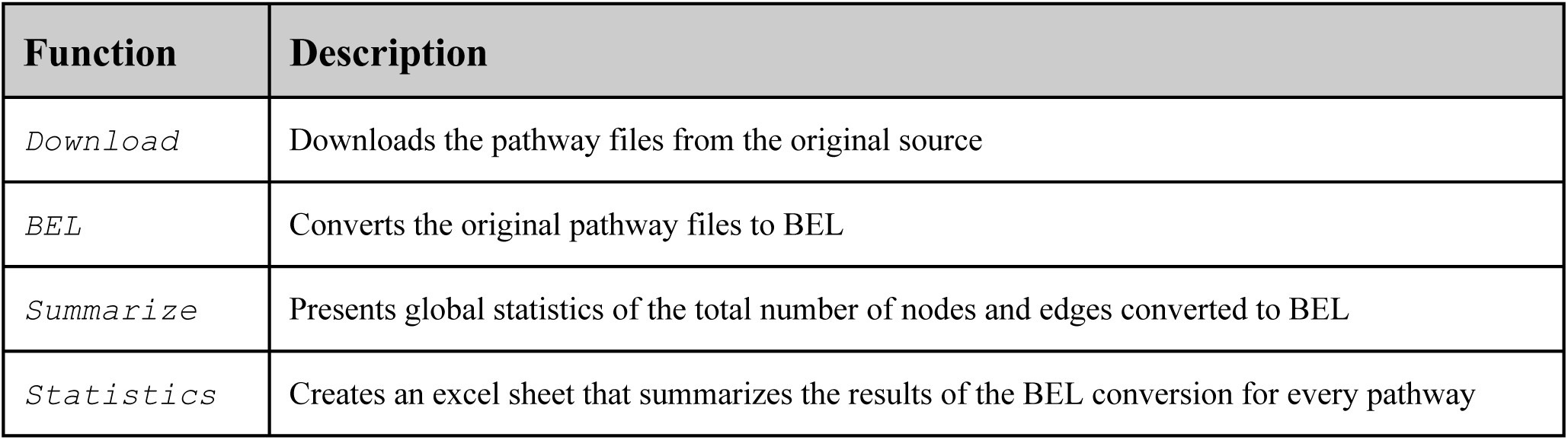
Core functions of the PathMe Python package.

### 3.2. PathMe Viewer

Beyond the software concerned with the integration of pathway knowledge, a novel web application (i.e. PathMe Viewer) was implemented for intuitive querying, browsing, and navigating of the normalized BEL networks. Queries can be submitted for a single or a set of pathways on the main page of the viewer, as illustrated in **Figure 2a**. The result of the query leads to a visualization, as seen in **Figure 2b**, that renders the corresponding network.

**Figure 2.**
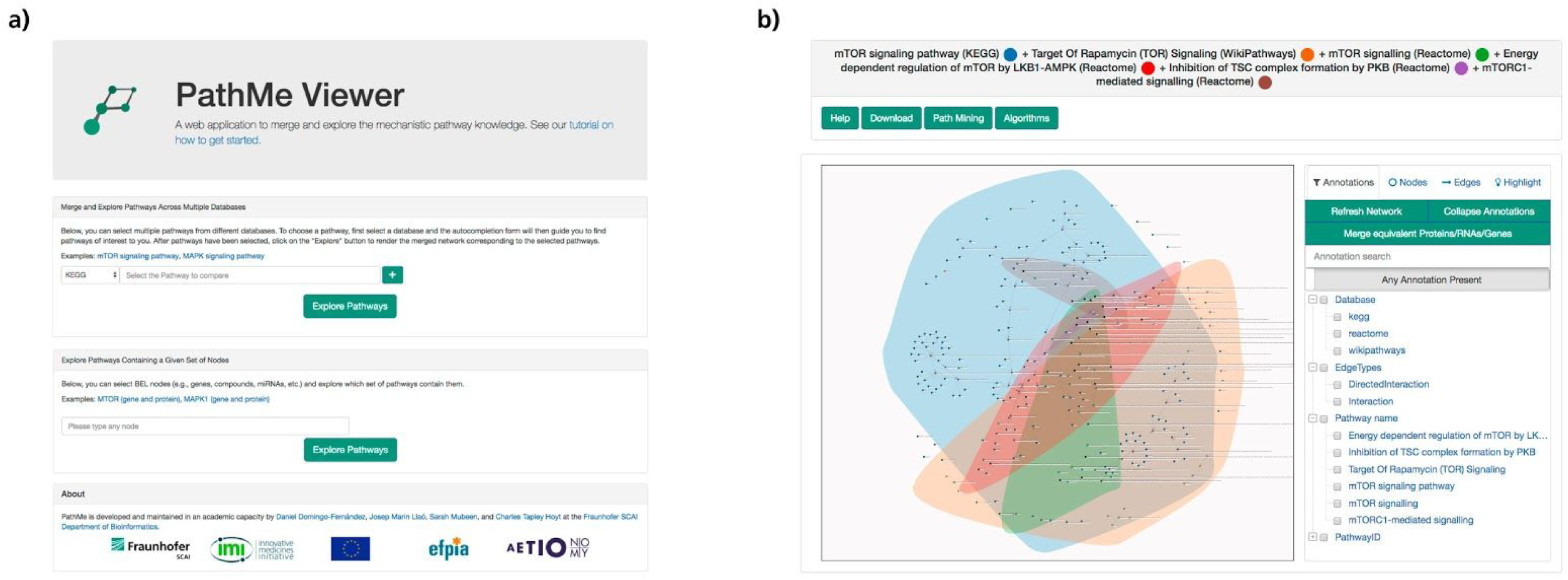
a) PathMe Viewer main page. b) The merged *mTOR signaling* network from KEGG, Reactome, and WikiPathways visualized in the PathMe Viewer. The highlighted regions mark the boundaries of each of the networks to definitively identify pathway landscapes, as defined by each of the network sources.

The PathMe Viewer is powered by multiple, built-in functionalities enabling users to navigate through the pathway(s). Although the initial network layout is defined by the D3 force algorithm which enables users to get a comprehensive overview of the relevant parts of the network, the network arrangement can also be customized by dragging and moving nodes around the viewer. Furthermore, node and edge meta-information can be accessed via double click. For nodes, this includes specifications on their name, function and namespace, while for edges, the pathway name, identifier and source database are provided. When multiple pathways are queried, marked boundaries delineate the topological landscape of each of the networks which synergistically contribute to the consolidated one to facilitate the exploration of pathway crosstalks (i.e. the interaction of pathways through their sharing of common entities) (**Figure 2b**). Furthermore, search and mining tools enable navigation of the resulting network such as selecting and filtering nodes/edges or calculating paths. Another novel feature of the viewer is the automatic identification of contradictory and consensus relationships across pathways (i.e., edges between identical nodes with equivalent or opposite relationship), which are highlighted in blue/red in the network. The viewer also incorporates a functionality which collapses all BEL proteins, RNA species and genes into gene nodes. This function was included in the viewer because of the interchangeable usage of these entities by the various databases which would both preclude the ability to fairly establish if there is overlap in the network topology and to conduct fair comparisons. Finally, network algorithms such as betweenness centrality can be used to quickly identify central nodes in the network or to calculate pathway similarity as we will present in the case scenario.

### 3.3. Software development techniques

Successful contributions to the bioinformatics domain are predicated by their ability to be replicated and reused. In line with community standards designed to foster these attributes, the PathMe and PathMe-Viewer packages use git for version control on GitHub, *flake8* to enforce code quality, *setuptools* to build distributions, *pyroma* to enforce package metadata standards, *sphinx* to build documentation, Read the Docs to host documentation, *py.test* as a unit and integration testing harness, and Travis-CI as a continuous integration server to run each of these with each commit (https://travis-ci.com/PathwayMerger/PathMe and https://travis-ci.com/PathwayMerger/PathMe-Viewer). Each package is distributed publicly through PyPI such that they can be included in other Python projects with requirements.txt or included in the setup.py using the *install_requires* setting without the need for complicated build steps or any other user configuration.

Because PathMe works on frequently updated external pathway data from multiple sources, it must be re-run frequently to incorporate those updates. Following the recommendation from Kim *et al.* (2018) for building reproducible environments for bioinformatics, we have encapsulated the entire PathMe workflow of acquiring, parsing, mapping, and normalizing the pathway resources within a Docker container such that it can be run on a *cron job* (i.e. a task scheduled to be re-run periodically). After, these changes are incorporated into the publicly available instance of the PathMe-Viewer. The cron job has the additional benefit that it reports when the formats of the underlying data change (which happens with moderate frequency) so the relevant PathMe components can be adapted. Also following the recommendation from Kim *et al.* for the scientific aspect of reproducibility, the three application scenarios presented in the next sections were conducted in IPython notebooks that are available and documented on GitHub (https://github.com/PathwayMerger/PathMe-Resources) that illustrate useful commands that might serve to assist similar future analyses.

### 3.4. Case scenario I: Global entity comparison across pathway databases

As a first application, we conducted a global comparison of biological entities across major modalities **(Figure 3)**. We would like to note that in order to ensure the quality of the comparison presented in this case scenario, this analysis exclusively uses a highly cited and peer-reviewed pathway set provided by WikiPathways (approximately 510) that has been approved and tagged for usability in data analysis. While we attempted to maximize the retention of biological entities, we found severe differences in the level of overlap across resources which demonstrates the importance of database integration to gain a holistic picture of pathway knowledge.

**Figure 3.**
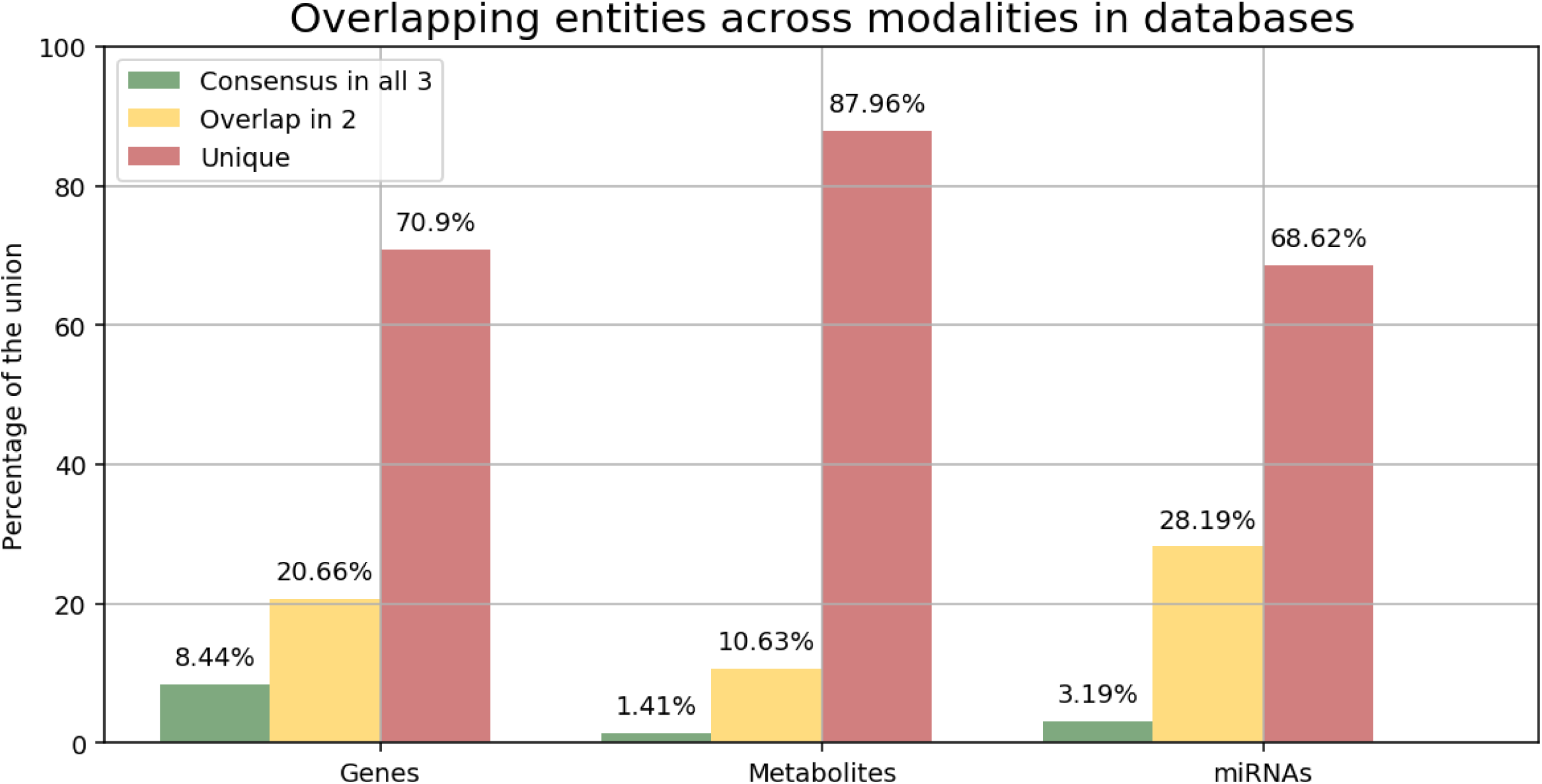
Overlapping entities across modalities in the three databases studied (i.e., KEGG, Reactome, and WikiPathways). The comparison analysis studied the degree of overlap for three different biological entities (i.e., genes, metabolites, and miRNAs) to evaluate whether entities are shared across databases (i.e., the ratio of the number of nodes present in all three databases to the number of nodes in the union of all databases for that modality), partially overlap (i.e., the ratio of the number of nodes present in only two databases to the number of nodes in the union of all databases for that modality) or are exclusive (i.e., the ratio of the number of nodes unique in one database to the number of nodes in the union of all databases for that modality). The classification of entities by their corresponding modalities are described in **Table 2.** We would like to remark that the analysis accounted for every entity present in the full set of pathways from the studied databases.

The degree of consensus of biological entities across all three databases was found to be relatively low, albeit variable across the assessed modalities **(Figure 3)**. The proportion of genes present in all databases was lower than the results obtained by Stobbe *et al.* (15%), though they exclusively focused their work on a set of metabolic pathways present in five major databases which included KEGG and Reactome. Total consensus of miRNAs in all three databases was unsurprisingly low due to a disproportionate representation of miRNA species across the databases. Specifically, as few as 13 miRNAs were derived from Reactome while 149 were present in KEGG. Similarly, the total consensus for metabolites was grossly deficient at less than 2%.

**Table 2.** Pathway database content statistics. Each cell reflects the unique number of entities for a given modality in its corresponding database. The *genes* modality comprises genes, mRNAs, and gene products as well as any modifications on those. The *metabolites* modality comprises biological entities from small molecules to cellular components. The *miRNAs* modality contains microRNA molecules. Finally, nodes that correspond to other pathways, molecular events, or biological processes (e.g., Gene Ontology (GO; Carbon *et al.*, 2017) terms) are included in the *biological processes* modality. The statistics reflect the status of the content available from KEGG and WikiPathways from the 13th of March, 2019 and the latest Reactome release (version 67) from the 13th of December, 2018.

| Modality | KEGG | Reactome | WikiPathways |
| --- | --- | --- | --- |
| Genes | 7289 | 8653 | 3361 |
| Metabolites | 4048 | 2712 | 655 |
| miRNAs | 149 | 13 | 91 |
| Biological processes | 418 | 2219 | 138 |

For partial overlap, we found that results varied across the three modalities, with a higher degree of overlap between miRNA species at approximately 30%, followed by genes with nearly 20%, and metabolites with approximately 11%. Accordingly, we found the proportion of distinct entities to be substantially higher than those present in any two or all three databases. The particularly low levels of overlap observed in all modalities can be largely attributed to several factors:

1. **The number of entities per modality across databases is highly variable (Table 2)**. Per definition, sets with significant variations in cardinality (i.e., set size) limit the likelihood of consensus since only a portion of the larger sets can intersect with the smaller ones. For instance, since KEGG contains 4048 metabolites and WikiPathways 655, the maximum consensus is constrained to the metabolites contained in WikiPathways (in this case, the maximum overlap would be 16.18%).
2. **The scope of the pathways comprised in each database varies.** Each database places a distinct emphasis on discrete aspects or regions of biological pathways which tend to be defined subjectively in the absence of standard nomenclatures, as outlined by (Domingo-Fernández, *et al.* 2019) who reported only 21 equivalent pathways between the three databases. Therefore, despite the presence of key biological players in all three databases, the majority of biological entities are particular to a single database. For example, over 200 glycan molecules are present in KEGG since this resource contains multiple pathways related to glycan metabolism (i.e., ‘Glycan biosynthesis and metabolism’) while they are absent in the others.
3. **Highly specific entity identifiers impede entity mappings with major standard nomenclatures.** Some entity identifiers have no discernible mapping to major standard nomenclatures because they exhibit a high degree of specificity. This is particularly evident for genes where curators have also captured entities such as specific protein events (e.g., “p-y641-stat6”), protein mutations (e.g., “activated fgfr1 mutants”), protein families (e.g., “lim kinases”), protein fragments (e.g., “ub c-terminal hh fragments”), or splicing events (e.g., “xbp1 mrna (spliced)”). A way to account for these high granular cases would be to standardize protein family names with resources such as Pfam (Finn *et al.*, 2015) or FamPlex (Bachman *et al*., 2018). For cases such as protein fragments or events, BEL enables their harmonization as they can be incorporated into its syntax (e.g., BEL *proteinModification(), fragment(), variant()*, etc.).
4. **Biological modalities can be broadly defined.** We characterize modalities to correspond to BEL node classes **(Table 2)**. For instance, the *genes* modality comprises gene, protein, and RNA BEL nodes. While this modality is clearly defined, there is a higher degree of variability in the entity types that can be that can be classified with the *metabolites* modality since the latter comprises a broad range of abundance BEL nodes (i.e., small molecules, cellular components, clinical measurements, or categories that do not fit in other BEL node classes; Pham *et al.*, 2019). Without the use of standard nomenclatures by the source databases, an extensive manual effort would be required to partition these modalities into more granular classifications. For example, the usage of GO as opposed to internally-defined terminologies to define cellular components would enable the categorization of cellular components into their own distinct modality. Similarly, the *biological processes* modality exhibits minimal overlap due to a lack of usage of standardized ontologies such as GO **(Supplementary Text)**.

### 3.5. Case scenario II: Comparing equivalent pathways in the three databases

Merging pathway knowledge enables analyzing the crosstalks for any set of pathways through the PathMe Viewer. As a case scenario, we used PathMe in conjunction with the viewer to explore the knowledge consolidated from 21 equivalent pathways across the three databases previously curated by Domingo-Fernández *et al.* **(Table 3)**. While conducting a cross-database pathway comparison previously required either extensive manual curation or harmonization of both entity identifiers and data formats on a case by case basis, this example illustrates how PathMe can be exploited to enable a systematic comparison of equivalent pathways.

**Table 3.** Consolidated pathway representations, their similarity indexes, and links to visualize the merged networks in the PathMe Viewer. A detailed analysis with the scripts to replicate the results and comments on the identified overlaps for each of the 21 equivalent pathways is available at https://nbviewer.jupyter.org/github/PathwayMerger/PathMe-Resources/blob/master/notebooks/case_scenarios/evaluating_similarity_equivalent_pathways.ipynb.

| KEGG | Reactome | WikiPathways | Pathway Similarity |
| --- | --- | --- | --- |

**Table 3.** Consolidated pathway representations, their similarity indexes, and links to visualize the merged networks in the PathMe Viewer.
|  |  |  | Index |
| --- | --- | --- | --- |
| Cell cycle | Cell Cycle | Cell Cycle | 0.70 |
| Toll-like receptor signaling pathway | Toll-Like Receptors Cascades | Toll-like Receptor Signaling Pathway | 0.62 |
| mTOR signaling pathway | mTOR signalling | Target Of Rapamycin (TOR) Signaling | 0.58 |
| Hedgehog signaling pathway | Signaling by Hedgehog | Hedgehog Signaling Pathway | 0.56 |
| Apoptosis | Apoptosis | Apoptosis | 0.43 |
| IL-17 signaling pathway | Interleukin-17 signaling | IL17 signaling pathway | 0.42 |
| PI3K-Akt signaling pathway | PI3K/AKT activation | PI3K-Akt Signaling Pathway | 0.42 |
| Wnt signaling pathway | Signaling by WNT | Wnt Signaling Pathway | 0.41 |
| MAPK signaling pathway | MAPK family signaling cascades | MAPK Signaling Pathway | 0.40 |
| B cell receptor signaling pathway | B Cell Receptor Signaling Pathway | Signaling by the B Cell Receptor (BCR) | 0.37 |
| Pentose phosphate pathway | Pentose phosphate pathway (hexose monophosphate shunt) | Pentose Phosphate Pathway | 0.33 |
| Citrate cycle (TCA cycle) | Citric acid cycle (TCA cycle) | TCA Cycle | 0.33 |
| Synthesis and degradation of ketone bodies | Ketone body metabolism | Synthesis and Degradation of Ketone Bodies | 0.33 |
| Notch signaling pathway | Signaling by NOTCH | Notch Signaling Pathway | 0.29 |
| DNA replication | DNA Replication | DNA Replication | 0.28 |
| Prolactin signaling pathway | Prolactin receptor signaling | Prolactin receptor signaling | 0.28 |
| TGF-beta signaling pathway | Signaling by TGF-beta family members | TGF-beta Signaling Pathway | 0.26 |
| Thyroid hormone synthesis | Thyroxine biosynthesis | Thyroxine (Thyroid Hormone) Production | 0.20 |
| Sphingolipid metabolism | Sphingolipid metabolism | Sphingolipid Metabolism | 0.16 |
| Mismatch repair | Mismatch Repair | Mismatch repair | 0.08 |
| Non-homologous end-joining | Nonhomologous End-Joining (NHEJ) | Non-homologous end joining | 0 |

To evaluate the degree of overlap between the three representations of each equivalent pathway, we used a variation of the Szymkiewicz–Simpson coefficient calculated for the common molecular nodes between the networks **(Equation 1)**.

Each of the 21 equivalent pathways showed partial overlap, except ‘Non-homologous end-joining’ which did not contain the pathway information required to convert the pathway into BEL in two of its original files. Among the equivalent pathways with the highest degree of similarity, we found well-studied pathways such as ‘Cell cycle’, ‘Toll-like receptor signaling’, ‘mTOR signaling’, Hedgehog signaling’, and ‘Apoptosis’. Although the three databases represent the most widely studied molecular players in each of these pathways, merging their knowledge assists in filling the gaps between the complex interactions occurring in these pathways. Pathways with low similarity, such as ‘TCA Cycle’ and ‘Sphingolipid Metabolism’, indicate the resources captured distinct aspects of the biology within the pathway. Unsurprisingly, this is in concordance with the findings reported by Stobbe *et al.* who conducted a comparison of the ‘TCA Cycle’ across five metabolic pathway databases. We would like to note that while previous approaches to characterize pathway similarity were purely gene-centric, our approach includes not only gene sets, but a range of modalities represented in pathways. Finally, beyond harmonizing entities and concepts, PathMe also harmonizes relationships, thus facilitating further analyses where pathway topology is included, as shown in the next case scenario.

### 3.6. Case scenario III: In-depth pathway analysis of mTOR signaling after superimposing its multiple representations

As a further application of the framework, we used the PathMe Viewer to conduct a detailed investigation of the mammalian target of rapamycin (mTOR) signaling pathway to demonstrate its utility in enriching pathway knowledge. In **Figure 4**, the consensus in terms of entity overlap across equivalent mTOR signaling pathways from each of the databases is depicted. All three databases are complementary to the others, but also possess some degree of overlap and thus are neither entirely identical nor distinct. Variability in the size of the mTOR signaling pathway, as measured by the number of nodes in each database, is also clearly discernible with KEGG contributing the largest proportion of distinct nodes to the heterogeneous, merged network **(Figure 4a)**.

**Figure 4.**
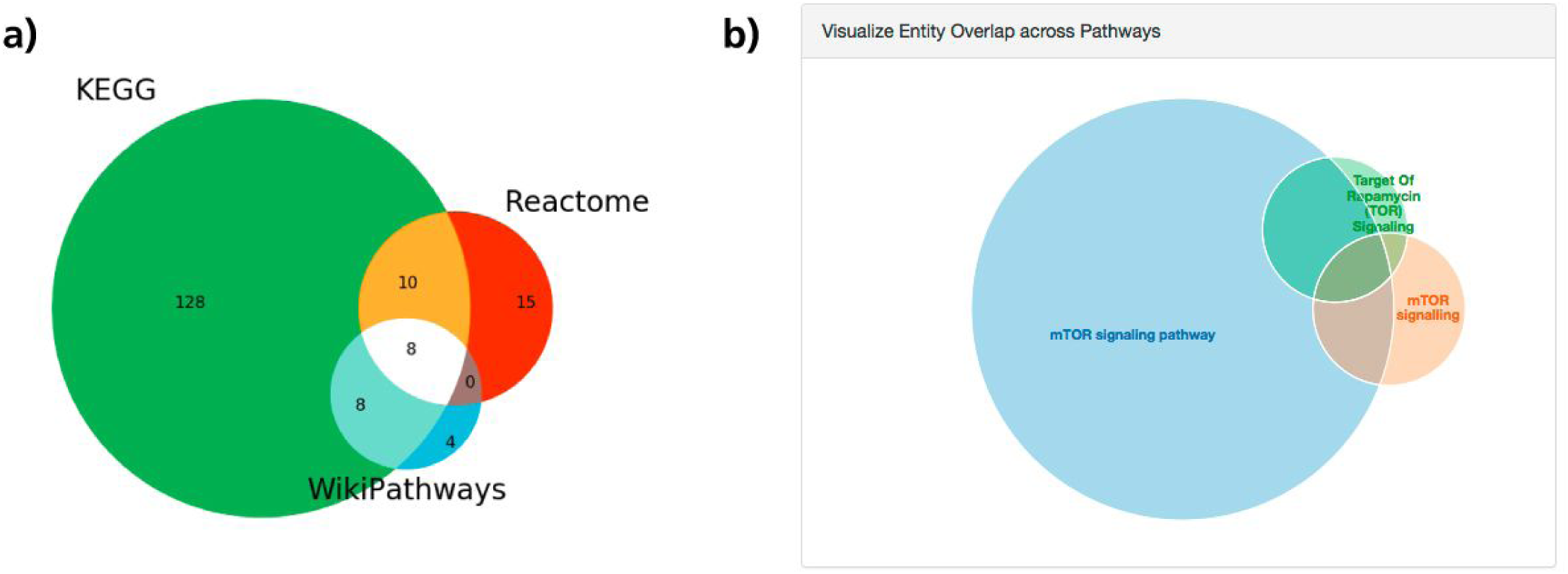
Node overlap between the three pathway representations of mTOR signaling. Note that this analysis includes all modalities harmonized by PathMe (e.g., genes, metabolites, miRNAs, biological processes). Particularly for mTOR signaling, while there are overlapping nodes between each pair of databases and between all three, each database also contributes unique nodes to the consolidated mTOR signaling network, providing a more comprehensive overview of mTOR signaling. In a), a more detailed analysis conducted on IPython notebooks shows the proportion of shared nodes across the three equivalent representations. Equivalent interactive Venn diagrams can also be generated directly from the PathMe Viewer for any set of pathways, as visualized in b).

A key functionality of PathMe Viewer is in the visualization and interactive exploration of pathways. In **Figure 5a and 5b**, using the viewer, an in-depth analysis of mTOR signaling reveals novel sets of interactions in the integrated network that are absent in individual mTOR signaling networks. The role of AKT signaling in modulating mTOR, as illustrated in **5a** and sourced from KEGG, has already been well-described in the literature (Altomare *et al.*, 2012; Memmott and Dennis, 2009). More notably, by superimposing the mTOR signaling pathway as defined in KEGG with its equivalent pathway from WikiPathways **(Figure 5b)**, an association between AKT1 and insulin related-processes becomes apparent in the merged network, though neither of the individual pathway sources connect the downstream effects of mTOR on insulin signaling. Nevertheless, the association between mTOR and insulin signaling through AKT modulation has been previously described in the literature (Le Bacquer *et al*., 2013). Additionally, bidirectional effects of mTOR have been demonstrated on AKT activity; these effects can vary both by the type of mTOR complex involved in the pathway and by negative feedback loops on insulin signaling, leading to altered states of AKT activity (Altomare *et al.*, 2012; Le Bacquer *et al*., 2013). As such, both pathways serve to complement each other in the integrated network, unraveling a connection which was hidden in disparate databases, though has been well-studied in the literature. Recent studies have also demonstrated that mTOR receives input from multiple pathways (Memmott and Dennis *et al*., 2009); a principal feature of the PathMe Viewer is in its capacity to directly visualize pathway crosstalk. While in the previous case scenario, crosstalk analyses were performed across equivalent pathways, in this case, using the viewer it would be possible to simultaneously visualize and explore different pathways which are evidenced to, or are possibly involved in, crosstalk with the mTOR signaling one.

**Figure 5.**
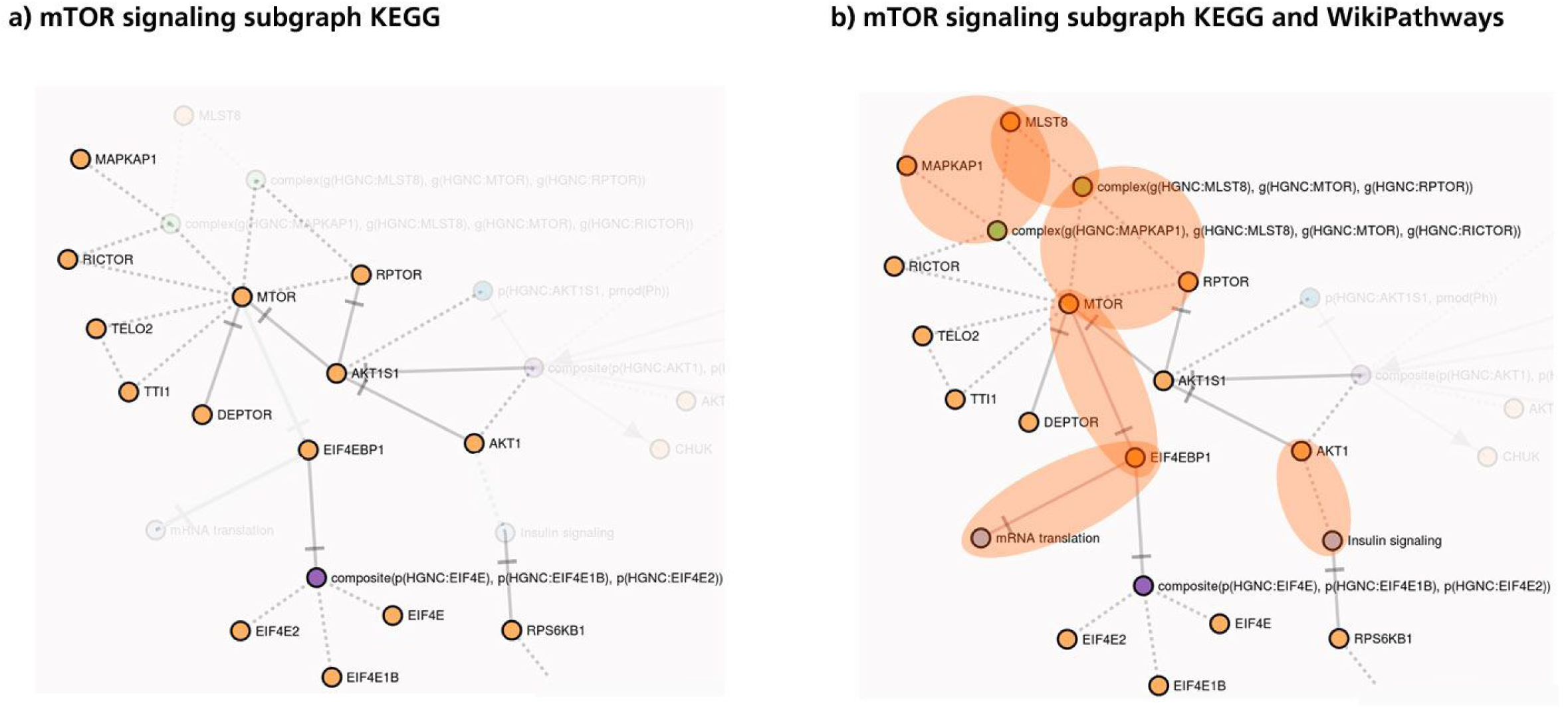
Superimposing mTOR signaling subgraphs from KEGG and WikiPathways. A deeper analysis performed using PathMe Viewer to visualize mTOR signaling subgraphs can be seen highlighting interactions present in a) KEGG only and b) KEGG and WikiPathways. The contribution of WikiPathways onto the pathway found in KEGG is highlighted in orange. Neither agreements nor discrepancies in topology are noted between these subgraphs though there are overlapping genes (e.g., mTOR, RICTOR, EIF4EBP1, etc.). Instead, subgraph a) is complemented by additional interactions superimposed onto the network, as visualized in subgraph b).

Similarly, by superimposing the mTOR signaling pathways from KEGG and WikiPathways, downstream interactions between mTOR and mRNA translation via EIF4EBP1 are evident **(Figure 5a and 5b)**. The inhibition of mTOR has been noted to be a potent repressor of protein translation while mTOR activation can stimulate mRNA translation through EIF4EBP1 (Gingras *et al.*, 2001; Choo *et al.*, 2008). Though only the inhibition relationship is captured in the viewer, in the presence of an activation relationship, the viewer also offers a feature to detect contradictory edges (e.g., node A increases node B in one pathway and decreases B in another) between identical nodes across two databases.

## 4. Conclusions

Parallel developments of pathway databases during recent decades have resulted in different formalization schemas, hampering the interoperability between these resources and creating data silos. Overcoming this obstacle is instrumental to better understand the mechanisms underlying pathway knowledge. Additionally, while our approach can accommodate multi-scale pathway information from divergent database formats into a singular and standardized schema, a minority of entities and interactions have no discernible equivalencies in BEL and, as such, had to be omitted. For instance, so far PathMe parsers can extract information from both humans and other species; however, despite the capacity of PathMe to harmonize human identifiers, additional work is required for the harmonization of identifiers belonging to other species as integration can help in identifying evolutionarily conserved genes and processes.

Here, we have presented a framework through which content across multiple pathway databases can be integrated and transformed into a unified schema. Although PathMe currently only incorporates content from three major pathway databases, its flexibility allows for future inclusion of additional pathway databases. Moreover, it holds the capacity to update its content and track developments in pathway knowledge, an issue earlier outlined by Wadi *et al*.. Finally, the three case scenarios presented illustrate how the framework can be used to assist researchers in addressing biological questions at varying degrees of specificity such as: i) integrating the pathway landscape at the database level, ii) comparing the degree of consensus at the pathway level, and iii) exploring pathway crosstalk and studying consensus at the molecular level.

Ultimately, we have shown how integrating pathway databases and making them interoperable enables global pathway representations that can contribute to a more holistic overview of pathway knowledge than the knowledge contained in any single one of the databases. In the future, these global representations could be used to conduct more comprehensive pathway-centric analyses. Furthermore, the reproducibility of previous pathway enrichment analyses could also be evaluated by replicating them using any database combination. In other words, what would happen if, instead of KEGG, an identical analysis were to be performed using the Reactome or Wikipathways databases, or any combination of the three?

## 5. Availability and requirements

**Project name:** PathMe

**Project home page:** https://github.com/PathwayMerger

**Operating system(s)**: Platform independent

**Programming language**: Python and JavaScript

**Other Requirements**: Python 3

**License**: Apache License 2.0

**Any restrictions to use by non-academics**: KEGG Commercial License

## List of abbreviations

BEL: : Biological Expression Language
BioPAX: : Biological Pathway Exchange
DSL: : Domain Specific Language
GO: : Gene Ontology
MVC: : Model-View-Controller
PyPI: : Python Package Index
RDF: : Resource Description Framework
SBGN: : Systems Biology Graphical Notation
SBML: : Systems Biology Markup Language
SPIA: : Signaling Pathway Impact Analysis
XML: : Extensible Markup Language

## Declarations

### Ethics approval and consent to participate

Not applicable

### Consent for publication

Not applicable

### Availability of data and materials

The datasets generated and/or analysed during the current study are available in the ComPath’s GitHub repository, [https://github.com/ComPath/resources]. The datasets generated and/or analysed during the current study are publicly available at https://github.com/PathwayMerger/PathMe-Resources.

### Authors’ contributions

DDF conceived and designed the study. SM and JML implemented the individual database parsers with help and supervision from DDF. CTH supervised harmonization into BEL using the PyBEL framework. DDF implemented the web application and conducted the application scenario with the help of SM. DDF, SM, and CTH wrote the paper. MHA participated in the critical definition of the concept, proposed and participated in drafting and commenting critically the manuscript.

## Acknowledgements

We are very grateful to the curators of KEGG, Reactome and WikiPathways for generating the raw content which was used in this work. Furthermore, we would like to thank Dr. Egon Willighagen for his helpful suggestions regarding WikiPathways data.

