## Supplementary Material for "PathMe: Merging and exploring mechanistic pathway knowledge"

### Supplement

#### Outline

- **KEGG to BEL:** Transformation process from KGML to BEL
- **Reactome to BEL**: Transformation process from RDF to BEL
- **WikiPathways to BEL**: Transformation process from RDF to BEL
- **BEL harmonization:** Summary of the harmonization process
- **Application scenario I:** semi-automatic term normalization
- **Application scenario II:** Exclusion of group nodes from analysis
- **Implementation details**
- **Integration in the ComPath ecosystem:** Integration of inter-database pathway mappings

### KEGG to BEL

The KEGG database (Kanehisa *et al.,* 2016) provides a custom, XML based exchange format for each pathway map contained in the database, known as KEGG Markup Language, or KGML. KEGG pathway maps are drawn and updated manually and KGML facilitates the representation of these pathways as graph objects, with entries corresponding to entity nodes and relations and reactions as edges between them.

These KGML files were accessed through the KEGG REST API and subsequently parsed by Pythons' element tree module. The resultant element tree was then traversed for KEGG entry elements to populate corresponding entity type dictionaries. Of the entity types represented in KEGG pathway maps, a subset of types relevant to our purposes were extracted. Identifiers for KEGG entities were prioritized as summarized in *Table S1*. Where mappings were not readily available, custom KEGG identifiers were retained.

| **KEGG Identifier** | **Harmonized Identifier Priorities** |
|---|---|
| Custom KEGG ID for human genes (prefixed hsa) | 1. HGNC<br>2. UniProt<br>3. KEGG custom gene (no available mapping) |
| Custom KEGG ID for compounds (prefixed cpd) | 1. ChEBI<br>2. PubChem<br>3. KEGG custom compound (no available mapping) |
| Custom KEGG ID for biological processes (prefixed map) | KEGG custom biological process |
| Custom KEGG ID for orthologs | KEGG custom ortholog |

**Table S1.** Harmoning KEGG identifiers.

The selected subset included those entity types which could be readily transformed into BEL nodes. The mapping of KEGG entities to their equivalent BEL nodes is summarized in *Table S2*.

| **KEGG Node** | **Equivalent in BEL** | **Explanation** |
|---|---|---|
| Gene/Enzyme | proteinAbundance(x) | Gene, enzyme or protein |
| Group | complexAbundance(x) | Complex of gene products |
| Compound | abundance(x) | Chemical compound |
| Map | bioprocess(x) | Pathway node |
| Reaction | reaction(reactants(x)), products(y)) | Reaction node |

**Table S2.** Mappings between nodes in KGML to BEL v2.0.

The representation of genes in KGML can be by way of single entities or groups of entities which are either similar, of the same family or those which are grouped together because of the ambiguity concerning the role of the genes. Similarly, KEGG compounds may also be represented as single or grouped entities, presumably contingent on their degree of similarity. KEGG genes, compounds and groups representing a complex of gene products were processed into BEL equivalents by optionally constructing BEL composites consisting of similar elements as defined by single identifiers in KGML files or flattened lists of similar entities.

Entities present in KEGG but with no clear BEL equivalence include KEGG hierarchies (i.e. BRITE) and unclassified types termed other. Additionally, KEGG orthologs also remain to be translated into BEL equivalents as we focus here exclusively on human pathways. However, in future we do intend to incorporate orthologs into our framework, hence KEGG orthologs are currently retrieved by our parser.

Similarly, we traversed the element tree for interaction types and extracted those which could be readily mapped to BEL edges. This amounted to the KEGG to BEL equivalencies outlined in *Table S3*. Those KEGG interactions which were not transformed into BEL edges because of ambiguity in translation or due to the absence of a BEL equivalent included state change, dissociation, missing interaction and hidden compound.

A significant portion of KEGG pathway nodes and edges have been directly captured with BEL. Those aforementioned types which we do not directly map to BEL are minimally represented in KEGG pathways, thus, though this is a source of information loss, this loss is mostly negligible. *Figure S1* summarizes the overall statistics for KEGG pathways under two conditions; in the first, unflattened condition, KGML nodes which are represented as grouped entities are translated into BEL composite abundances. In the second, flattened condition, KGML nodes represented as groups are not combined in their translation into BEL and are represented solely as individual entities. Thus, the number of BEL nodes in the unflattened condition is greater than that of the flattened condition by way of the inclusion of composites. In addition to the discrepancy between BEL nodes in the flattened versus unflattened condition, the number of BEL nodes is notably less than the number of entities in the XML file *(Figure S1)*. Though a minority of those entities that were not translated from XML into BEL include orthologs, the majority of ostensible information loss is in genes and compounds. While entries in KGML files can be repeated, with the possibility of multiple, identical entries with unique IDs, all nodes are unique in BEL graph representations. Thus, this discrepancy in the number of entities can be largely attributed to the removal of duplicates.

| **KEGG Edge** | **Equivalent in BEL** | **Explanation** |
|---|---|---|
| Activation | x increases activity(y) | Subject increases the activity of object |
| Inhibition | x decreases activity(y) | Subject decreases the activity of object |
| Expression | x increases rnaAbundance(y) | Subject increases expression object |
| Repression | x decreases rnaAbundance(y) | Subject decreases expression object |
| Phosphorylation | activity(x) increases proteinAbundance(y, proteinModification(Ph)) | Subject increases phosphorylation of object |
| Dephosphorylation | activity(x) decreases proteinAbundance(y, proteinModification(Ph)) | Subject decreases phosphorylation of object |
| Glycosylation | activity(x) increases proteinAbundance(y, proteinModification(Glyco)) | Subject increases glycosylation object |
| Ubiquitination | activity(x) increases proteinAbundance(y, proteinModification(Ub)) | Subject increases ubiquitination object |
| Methylation | activity(x) increases proteinAbundance(y, proteinModification(Me)) | Subject increases methylation object |
| Indirect effect | x association y | Subject affects object but details are not given |
| Compound | x association y | Association event |

| | | |
|---|---|---|
| Binding/association | x association y | Association event |
| Irreversible reaction | catalyticActivity(x) increases reaction(reactants(y), products(z)) | Uni-directional reaction |
| Reversible reaction | catalyticActivity(x) increases reaction(reactants(y), products(z)), catalyticActivity(x) increases reaction(reactants(z), products(y)) | Bi-directional reaction |

**Table S3.** Mappings between edges in KGML to BEL v2.0.

A noticeable difference in the number of edges can be seen in BEL relative to those present in KGML *(Figure S2)*. This difference can be attributed to the generation of additional edges including those edges which delineate membership of entities in complexes and those formed between reaction elements. A significantly more pronounced difference between BEL edges and KGML interactions in the flattened condition can also be seen in *Figure S2*. This is due to the generation of edges between all composite components and the neighbours of the composites while in the unflattened mode, edges are formed exclusively between composites and their neighbours.

A detailed look into the KGML to BEL statistics can be seen in the KEGG to BEL Jupyter notebook [https://nbviewer.jupyter.org/github/PathwayMerger/PathMe-Resources/blob/master/notebooks/kegg/kegg_to_bel_statistics.ipynb].

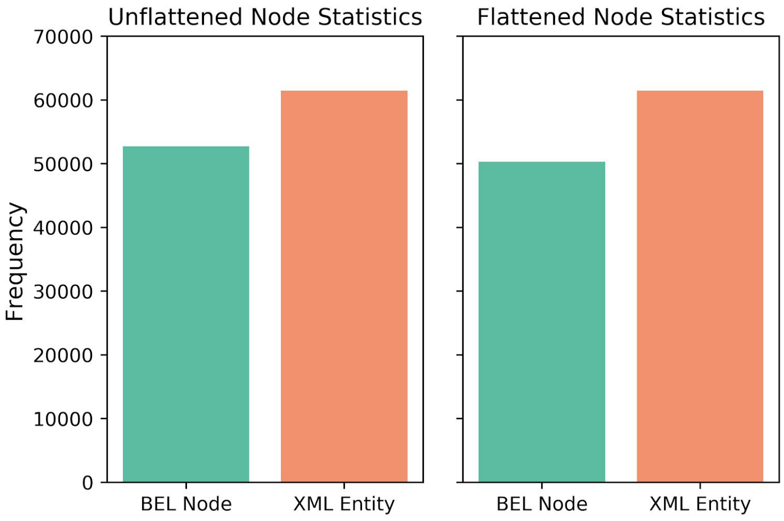


**Figure S1.** BEL and XML (KGML) node statistics in unflattened versus flattened conditions for all KEGG pathways. The difference in the number of BEL nodes in the unflattened versus flattened condition, where the value of the former is slightly larger, can be attributed to the inclusion of composites in the unflattened condition. The discrepancy in the number of nodes in BEL versus KGML can be largely attributed to the removal of duplicate nodes present in KGML files.

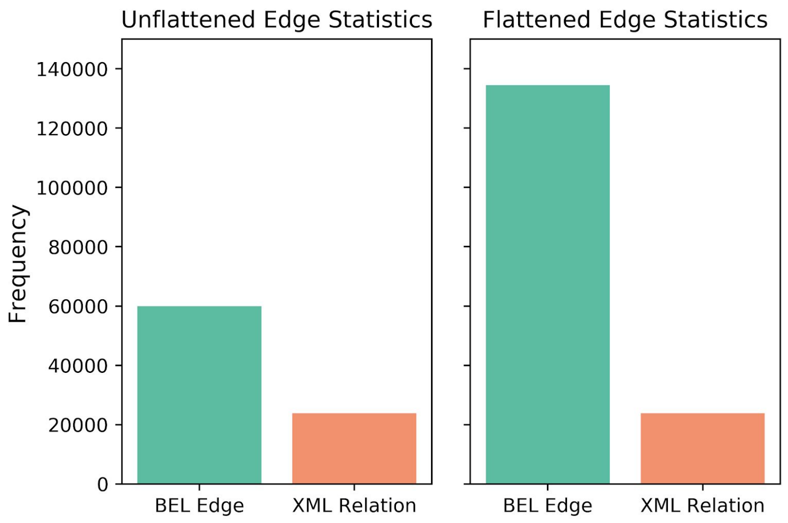


**Figure S2.** BEL and XML (KGML) edge statistics in unflattened versus flattened conditions for all KEGG pathways. The difference between the number of BEL edges in the unflattened condition is due to the generation of additional edges (ex. Designating membership in complexes, edges between reaction nodes and their reactants and reaction nodes and their products). The more pronounced difference between BEL edges and KGML interactions is partly due to this aforementioned generation of additional edges and largely due to the generation of edges between all composite *components* to each of the composites' neighbours. In contrast, in the un-flattened condition, edges are restricted to those between composites and their neighbours.

### Reactome to BEL

Pathways from the Reactome (Fabregat *et al.*, 2017) database can be downloaded in the PSI-MITAB, SBML, SBGN and BioPAX level 2 and 3 formats. Additionally, the database can be downloaded in Neo4j as an interconnected Reactome Graph.

We downloaded Reactome pathways in the BioPAX format through the European Bioinformatics Institutes' (EBI) File Transfer Protocol (FTP) which provides access to RDF datasets in bulk. Because BioPAX is defined using the standard OWL (RDF/XML) syntax, this format can also be used with RDF/OWL tools such as reasoners or triplestores. The RDF file for pathways in humans was then parsed using various SPARQL queries to extract those entity and interaction types which could be directly mapped to nodes and edges in BEL. Identifiers for entities in Reactome were prioritized as described in *Table S4*.

| **Reactome Identifier** | **Harmonized Identifier Priorities** |
|---|---|
| Reactome ID for genes/proteins | 1. HGNC<br>2. UniProt<br>3. Ensembl<br>4. Reactome custom gene/protein (no available mapping) |
| Reactome ID for compounds | 1. ChEBI<br>2. PubChem<br>3. Reactome custom compound (no available mapping) |

**Table S4.** Harmoning Reactome identifiers.

Mappings from each entity and relation type in Reactome to their equivalent BEL representations were then generated, as summarized in *Table S5.* The final transformation to BEL is done using the PyBEL package (Hoyt *et al*., 2018), where similarly to KEGG, each node/edge is translated to the PyBEL data structure in order to serialize Reactome to BEL. It is important to remark that RDF files were available for pathways from other species for which the parser can also be applied.

| **Reactome Node** | **Equivalent BEL Node** | **Explanation** |
|---|---|---|
| Protein | proteinAbundance(x) | Node is gene or protein |
| SmallMolecule | abundance(x) | Node can be any small molecule |
| Pathway | biologicalProcess(x) | Node is a pathway |
| DNA | geneAbundance(x) | Node is the abundance of the gene |
| RNA | rnaAbundance(x) | Node is the abundance of RNA |
| Complex | complexAbundance(x) | Node is a complex |
| **Reactome Edge** | **Equivalent BEL Edge** | **Explanation** |
| Activation | x increases activity(y) | Subject increases the activity of object |
| Inhibition | x decreases activity(y) | Subject decreases the activity of object |

**Table S5.** Mappings between nodes and edges in the BioPAX format to BEL v2.0.

The statistics of the conversion of Reactome entities and interactions into BEL nodes and edges are presented in the Reactome to BEL Jupyter notebook [https://nbviewer.jupyter.org/github/PathwayMerger/PathMe-Resources/blob/master/notebooks/reactome/reactome_to_bel_statistics.ipynb]. The results are summarized in *Figure S3*.

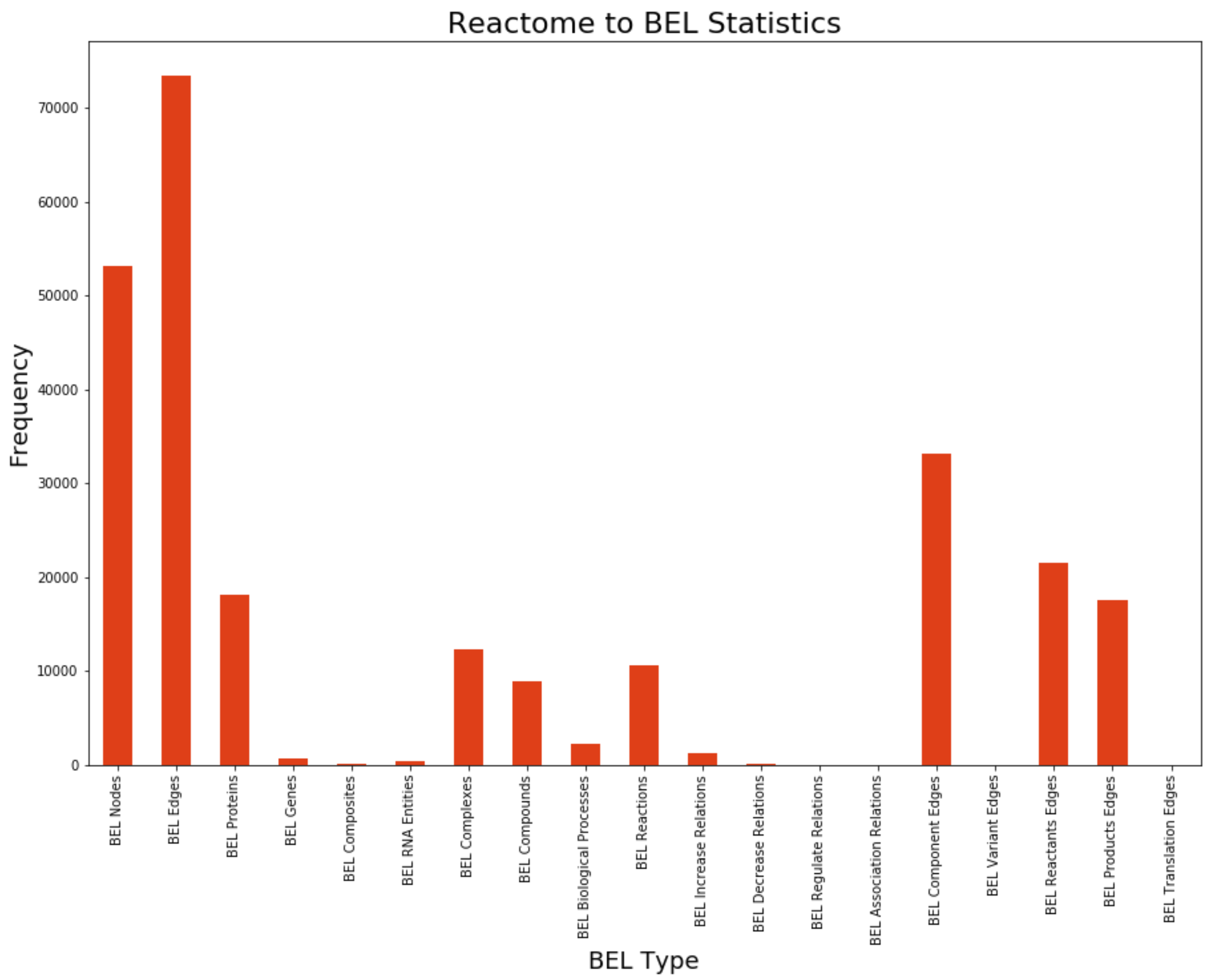


**Figure S3.** Statistics of the conversion of Reactome entities and interactions into BEL nodes and edges, respectively. All human pathways from the Reactome RDF file were considered.

### WikiPathways to BEL

WikiPathways (Slenter *et al*., 2018) provides a Semantic Web Portal where its content can be downloaded as RDF (Waagmeester *et al*., 2016). The RDF files contain pathway information that we parsed using multiple SPARQL queries in order to extract the meta-information corresponding to each node/edge in the RDF graph. At this point, we would like to remark that by default the package only parses the original WikiPathways pathways and not other imported pathways from other resources such as Reactome. Additionally, this package parses only human pathways, though pathways from other species can also be applied.

After we extracted and classified the data from RDF, identifiers for WikiPathways entities were prioritized as detailed in *Table S6*.

| **WikiPathways Identifier** | **Harmonized Identifier Priorities** |
|---|---|
| WikiPathways ID for genes/proteins | 1. HGNC<br>2. UniProt |

| | |
|---|---|
| | 3. Ensembl<br>4. WikiPathways custom gene/protein (no available mapping) |
| WikiPathways ID for compounds | 1. ChEBI<br>2. WikiPathways custom compound (no available mapping) |

**Table S6.** Harmoning WikiPathways identifiers.

To subsequently translate pathways in RDF format to BEL, we had to first reach a consensus between the types of nodes and edges present in WikiPathways and their equivalence in BEL. *Table S7* below shows the equivalences. The final transformation to BEL is done using the *PyBEL* package (Hoyt *et al*., 2018), where similarly to Reactome, each node/edge is translated to the PyBEL data structure in order to serialize WikiPathways to BEL.

| **WikiPathways RDF Node** | **Equivalent BEL Node** | **Explanation** |
|---|---|---|
| Protein/GeneProduct | proteinAbundance(x) | Node is a protein or gene product |
| DataNode | abundance(x) | Node can be abundance of any entity |
| Metabolite | | |
| Pathway | biologicalProcess(x) | Node is a pathway |
| Rna | rnaAbundance(x) | Node is abundance of RNA |
| Complex | complexAbundance(x,y) | Node is a complex |
| Conversion | reaction(reactants(x), products(y)) | Node is a reaction |
| **WikiPathways RDF Edge** | **Equivalent BEL Edge** | **Explanation** |
| Stimulation | x increases activity(y) | Subject increases the activity of object |
| Catalysis | x increases reaction(y) | Activity of subject increase transformation of reactants to products |
| Inhibition | x decreases activity(y) | Subject decreases the activity of object |
| DirectedInteraction | x association y | Subject has association with object |
| TranscriptionTranslation | x translatedTo y | RNA members translated to protein members |
| ComplexBinding | complexAbundance(x,y) | This information is duplicated with the Complex node, since a Complex can also be considered as an interaction |

**Table S7.** Equivalencies between WikiPathways RDF and BEL v2.0.

The statistics for the translation of WikiPathways entities and interactions into BEL nodes and edges can be found in the WikiPathways to BEL Jupyter notebook [https://nbviewer.jupyter.org/github/PathwayMerger/PathMe-Resources/blob/master/notebooks/wikipathways/wikipathways_to_bel_statistics.ipynb]. The results for the conversion of WikiPathways to BEL are summarized in *Figure S4*.

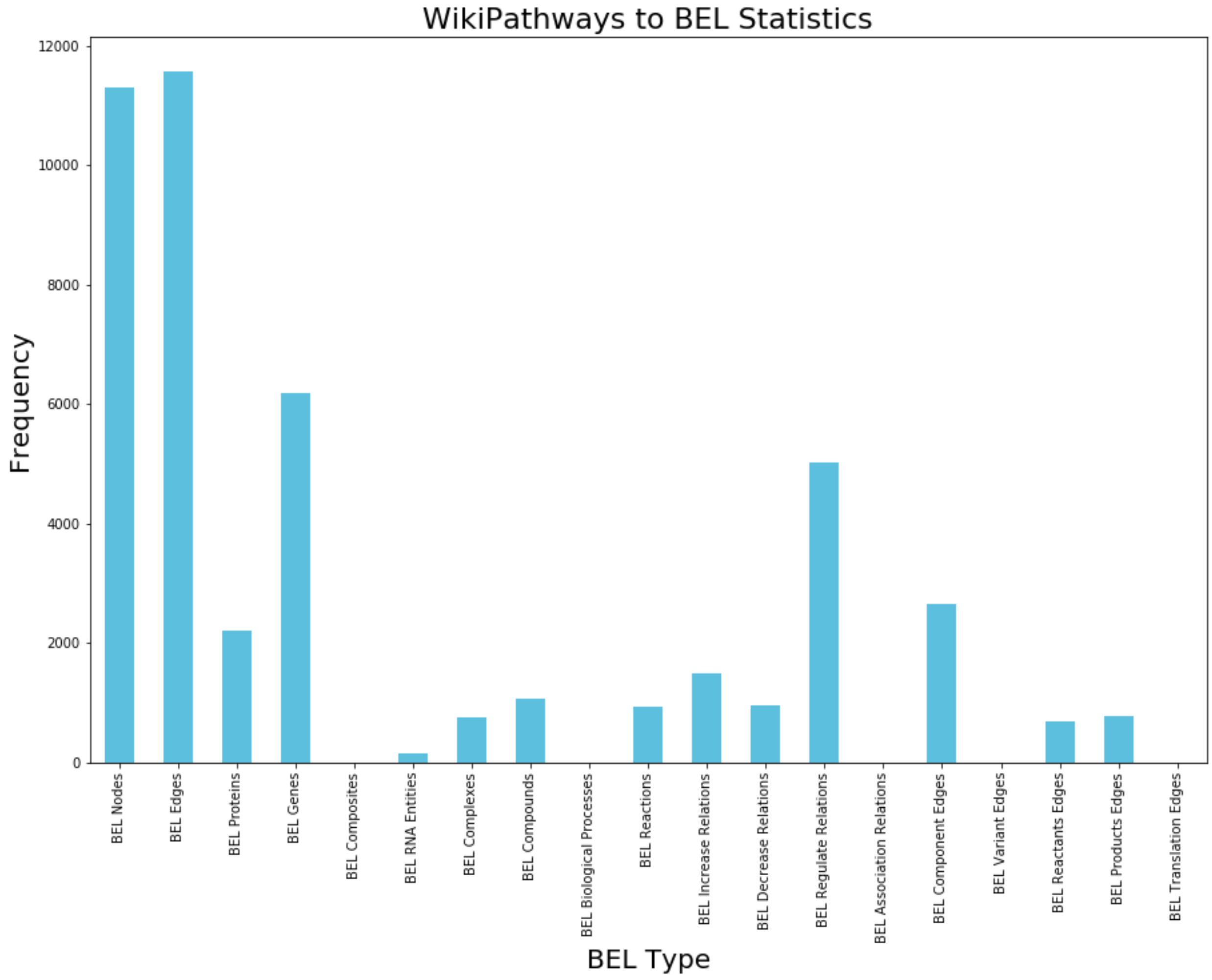


**Figure S4.** Statistics of the conversion of WikiPathways entities and interactions into BEL nodes and edges, respectively.

### BEL harmonization

As we have shown in the previous sections, BEL v2.0 is capable of capturing almost all of the information from the three different databases and thus, harmonizing the pathway knowledge in a fully interoperable, specified and structured schema. However, as Sales and colleagues discussed in their work, one can represent nodes containing multiple entities or group nodes (e.g., protein complexes, protein families, etc.), by one of two approaches *(Figure S5)*. The first approach represents a set of member entities as participants in a group, corresponding to a single node which can then be connected to its neighbours by its original edges. Conversely, the latter approach creates individual nodes for each of the member entities of a defined group. These member entities can then be connected with the original neighbours of the group node. While both approaches lead to a completely different network topology, they are both valid since they are suited for different applications.

The first formalism, which represents group nodes as individual nodes, is particularly well suited in cases where the node groups represent a protein complex. Whilst generating edges between member parts of a complex and its neighbours would alter the biological meaning of a relationship between a complex and its interacting entity, this first approach preserves the biological context, and thus, facilitates interpretation and visualization. On the other hand, the latter approach which segregates group node member entities into individual nodes, results in the generation of a comparatively large number of edges and is readily suitable for manipulating the network structure to facilitate working with graph algorithms (e.g., signaling and propagation) and for the representation of protein families, groups of similar proteins or where there is ambiguity concerning the role of the proteins involved in interactions.

Both of these approaches are accounted for in PathMe; by tuning the module with different settings, group nodes can be in the non-flattened *(Figure S5.b)* or flattened *(Figure S5.a)* condition, thus enabling a customized yet transparent pathway reconstruction in BEL.

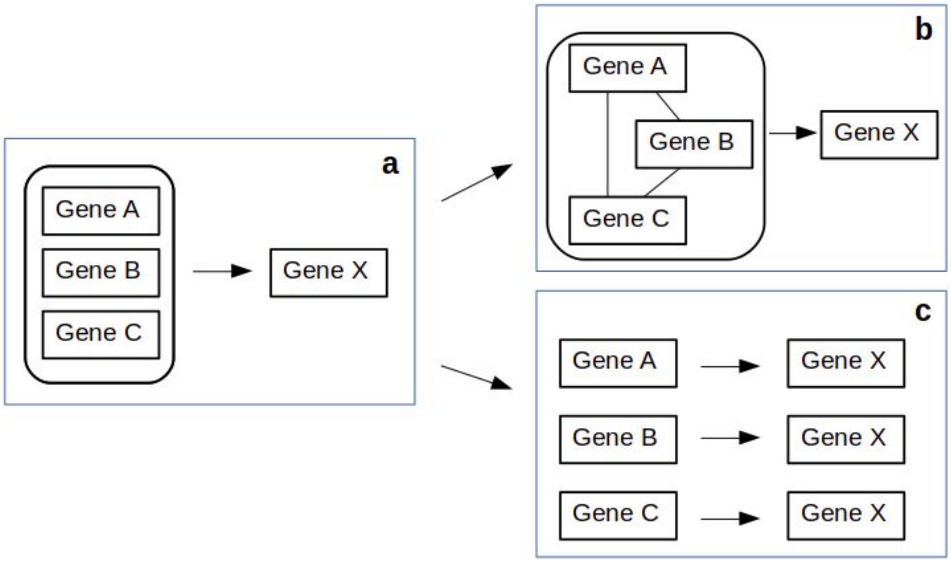


**Figure S5.** Group nodes representation approaches. Node groups, (**a**), can have edges between each member of the group and a single edge from the group to its neighbours (**b**), or edges between each member of the group to the groups' neighbours (**c**).

### Application scenario I: semi-automatic term normalization

| **Modality** | **Database** | **Rules applied in the normalization process** | **General comments** |
|---|---|---|---|
| Genes | KEGG | - | - |
| | WikiPathways | - | We found outdated EC identifiers and non-human proteins that could not be normalized. |
| | Reactome | Several individual identifiers correspond to multiple entities partaking in reactions. However, since curators did not use a | Numerous identifiers described protein families (e.g., "specific granule membrane proteins") or protein variations/modifications (e.g., "fgfr1 |

| | | | |
|---|---|---|---|
| | | consistent grammar to separate the entities (i.e., while some curators used commas to separate the terms, others used slashes), several rules had to be applied to extract the terms and conduct the normalization. | mutants with enhanced kinase activity“). Since these terms do not follow a standard nomenclature, they cannot be normalized and require manual curation. |
| Metabolites | KEGG | All entities were lower-cased to facilitate the normalization. | *KEGG:* Several internal identifiers are associated with glycan metabolism (e.g., gl:G00126) and thus, require special mappings to standard nomenclatures. *Reactome:* Proteins and cellular compartments are coded as metabolites. *WikiPathways:* Some genes were coded as metabolites and had to be manually moved to the *genes* modality. |
| | WikiPathways | | |
| | Reactome | | |
| miRNA | KEGG | - | - |
| | WikiPathways | - | |
| | Reactome | Necessary to capitalize the identifiers (e.g., from miR10 to MIR10) and remove prefixes, like “genes”, from the identifier. | |
| Biological Processes | KEGG | - | The majority of the terms used in this modality refer to pathways. The lack of a standardized pathway nomenclature, as demonstrated by Domingo-Fernández *et al.,* leads to their minimal overlap across databases (i.e., there are no overlaps except for the term “Apoptosis”). |
| | WikiPathways | - | |
| | Reactome | - | |

**Table S8.** Summary of the rules applied to normalize entities across databases for each modality.

### Application scenario II: exclusion of group nodes in the comparison analysis

The group nodes (i.e., composites, complexes, and reactions) were intentionally excluded when calculating the similarity index. There were two main reasons for doing so:

1. There are an unlimited number of possibilities when generating group nodes because of the combinatorial complexity of grouping members (i.e., a group node has nn possible combinations). This directly conflicts with the formalism of a set, which is by definition limited. Therefore, by exclusively using individual nodes in the pathway similarity calculations, the number of possible nodes are restricted to a finite number (i.e., the number of known/characterized molecular entities).
2. It is unlikely to find a match between two group nodes since all members in the group must be identical. This can distort the results when comparing a pathway to one having numerous group nodes versus one that possesses relatively few groups or none altogether. In such a case, the former would have a lower similarity score than the latter even if the group nodes are composed of components that are nearly identical.

### Implementations details

| Library | Purpose | Reference |
|---|---|---|
| RDFLib | Parsing and handling of RDF resources in WikiPathways and Reactome | https://pypi.org/project/rdflib/ |
| Python XML Module | Parsing and handling of KGML files in KEGG | https://docs.python.org/3/library/xml.html |
| PyBEL | Handling and creating BEL graphs | https://pypi.org/project/pybel/ |
| PyBEL-Tools | Enrichment of BEL graphs | https://pypi.org/project/pybel-tools/ |
| Pandas | Data handling, statistics generation | https://pypi.org/project/pandas/ |
| Bio2BEL KEGG | Query updated KEGG pathways | https://pypi.org/project/bio2bel-kegg/ |
| Bio2BEL Reactome | Query updated Reactome pathways | https://pypi.org/project/bio2bel-reactome/ |
| Bio2BEL WikiPathways | Query updated WikiPathways pathways | https://pypi.org/project/bio2bel-wikipathways/ |
| Bio2BEL HGNC | Mapping gene and protein identifiers | https://pypi.org/project/bio2bel-hgnc/ |
| Bio2BEL ChEBI | Mapping across metabolite identifiers | https://pypi.org/project/bio2bel-chebi/ |

**Table S9.** Python packages used for the harmonization of pathway database content into BEL.

| Technology | Functionality |
|---|---|
| MySQL | Relational database management system |
| Flask toolbox | An integrated web server and template manager that wraps many of the low level functions in an easy-to-manage programming interface |
| Docker | Reproducible deployment in any OS |

**Table S10**: A summary of back-end technologies used by the PathMe Viewer.

| Javascript Library | Functionality |
|---|---|
| jQuery | Provides manipulation of DOM, CSS, and general-purpose javascript |
| D3.js | Network visualization |
| InspireTree | Builds tree for annotation browser |

| | |
|---|---|
| Bootstrap Toggle | User interface for Toggle buttons |
| SVG.js | Export SVG images |

**Table S11**: A summary of front-end technologies used by the PathMe Viewer.

### Integration in the ComPath ecosystem

Even after databases have been harmonized into a common schema, one cannot directly explore agreements and pathway demarcations due to the lack of cross-references and mappings. As we point out in the main manuscript, there are several reasons that make automatically linking pathways from disparate database difficult, thus necessitating an exhaustive manual evaluation of the possible mappings for each database and for each pathway. Since the mappings for the three databases showcased in this paper were already generated (Domingo-Fernández *et al*., 2018), we demonstrate the utility of PathMe in the application presented in the main manuscript where we explore the crosstalks of equivalent pathways in the three databases. We would like to remark that this approach is only possible when pathways have been mapped and made fully interoperable. Therefore, the development of PathMe should be tightly related to the curation aspect of the ComPath project to generate more mappings in the future.

In order to facilitate the crosstalk between both web applications, we implemented the PathMe Viewer in a modularized manner so it can be deployed jointly with ComPath while maintaining its independence. While ComPath benefits by linking each pathway information page to the corresponding network using the PathMe Viewer, the latter can use the pathway mappings from ComPath.
